# ModSeqR: An R package for efficient analysis of modified nucleotide data

**DOI:** 10.1101/2025.11.10.687705

**Authors:** Hailey E. Zimmerman, Jordan L Moore, Ryan H. Miller, Isaac Stirland, Andrew Jenkins, Erin Saito, Tim Jenkins, Jonathon T. Hill

## Abstract

DNA methylation regulates a wide range of biological processes, including gene expression, disease progression, and cell identity. Long-read technologies now enable more comprehensive and accurate methylome analyses than ever before, but they are hindered by the computational resources needed to analyze the massive datasets. Here, we present the CH3 file format, which aids data storage and transfer by reducing file sizes by more than 95%, and the ModSeqR R package, which builds on the CH3 format and a database backend to enable a broad range of epigenetic analyses. Together, these tools enable high-throughput methylation analysis while minimizing computational resource requirements.

## Background

Although nearly every cell in an organism contains the same DNA sequence, DNA function varies widely across cell types, over developmental time, and in response to environmental stimuli. This regulatory diversity creates the rich mosaic of cellular behaviors that makes multicellular life possible. One of the most important mechanisms underlying this phenomenon is the chemical modification of DNA and RNA nucleotides^1^. These modifications encompass a wide range of chemical changes and are only beginning to be systematically described and understood. Thus, accurate detection and analysis of DNA modification patterns are essential for understanding development, disease mechanisms, and responses to environmental changes^2^.

The most well-characterized DNA modification is the addition of methyl groups to the fifth carbon of cytosine residues, typically within cytosine-guanine dinucleotide (CpG) contexts. DNA methylation plays vital roles in regulating cellular identity and function in humans and other species. Aberrant methylation patterns can destabilize the genome and disrupt normal gene expression, contributing to numerous diseases and serving as important diagnostic biomarkers^3,4,5^. Traditional approaches to DNA methylation profiling have relied heavily on microarray technologies such as the Illumina MethylationEpic Bead Chip. These arrays, while cost-effective and high-throughput, are inherently limited in genomic coverage and resolution^6^. Recent sequencing-based methods, including bisulfite sequencing and enzymatic methyl-seq (EM-seq), have extended our ability to interrogate methylation on a genomic scale. However, these technologies introduce additional limitations, including biases introduced during cytosine conversion and PCR amplification, as well as computational analysis challenges due to the number of mismatches introduced by the conversion process. In contrast, native methylation sequencing directly analyzes unamplified, unconverted DNA, avoiding these limitations. Platforms such as Oxford Nanopore Technologies (ONT) and PacBio have enabled native, PCR-free sequencing and subsequent genome-wide detection of DNA modifications in real time and at high resolution^7,8^. These third-generation technologies offer more comprehensive and unbiased epigenetic analyses than array-based platforms and other traditional methods^9^, thus providing significant technical advantages in most use cases, especially when studying genomes from non-model organisms or exploring genomic regions that are poorly represented by commercial arrays^10^.

Despite its promise, native modified-base sequencing data remains challenging to work with due to its large size and lack of streamlined analytical tools^11^. A single pass of the human genome (1x) produces 3-5 gigabases (Gbases) of data. Thus, sequencing at deeper coverages or with multiple samples can produce terabytes of information. Several tools have been developed to address this issue by compressing raw sequencing data^12^, but they fall short in providing support for further analysis. Other software packages provide differential methylation analysis capabilities, but were primarily developed for short-read bisulfite sequencing data^13^. Consequently, their performance is suboptimal for analyzing data from other paradigms, such as long-read sequencing or studying alternative DNA modifications^14^. As a result, there remains an urgent need for efficient tools to analyze downstream data from long-read, native DNA modification sequencing.

To address these gaps, we developed a two-part solution: a highly efficient modified-nucleotide data storage format (CH3) that reduces nanopore data sizes by 90-95%, and ModSeqR, a publicly available R package for fast, flexible downstream analysis. ModSeqR supports differential methylation analysis at single-base, sliding windows, or regional levels, includes built-in visualization and read-classification tools, and offers customizable parameters for diverse experimental designs. Here we present the package’s workflow, features, and performance.

## Results

### CH3 - Efficient DNA Modification Data Storage

Methylation sequencing generates large volumes of raw data, typically in the form of binary alignment map (bam) files containing raw read sequences. When modified base calling is enabled, two tags are added to each record in the file: one for the modified base call positions and a second with probabilities for modification at the position. In order to analyze the data, these files undergo a series of pre-processing steps, as summarized in **Figure 1a**. This process includes alignment, merging, sorting, and indexing. Finally, base modification predictions—such as methylation events—can be extracted and stored in .tsv files using tools like the modkit package (https://github.com/nanoporetech/modkit). While convenient as a human-readable format, these plain-text call files are often unmanageably large, presenting challenges for downstream analysis, storage, and sharing.

**Figure 1:**
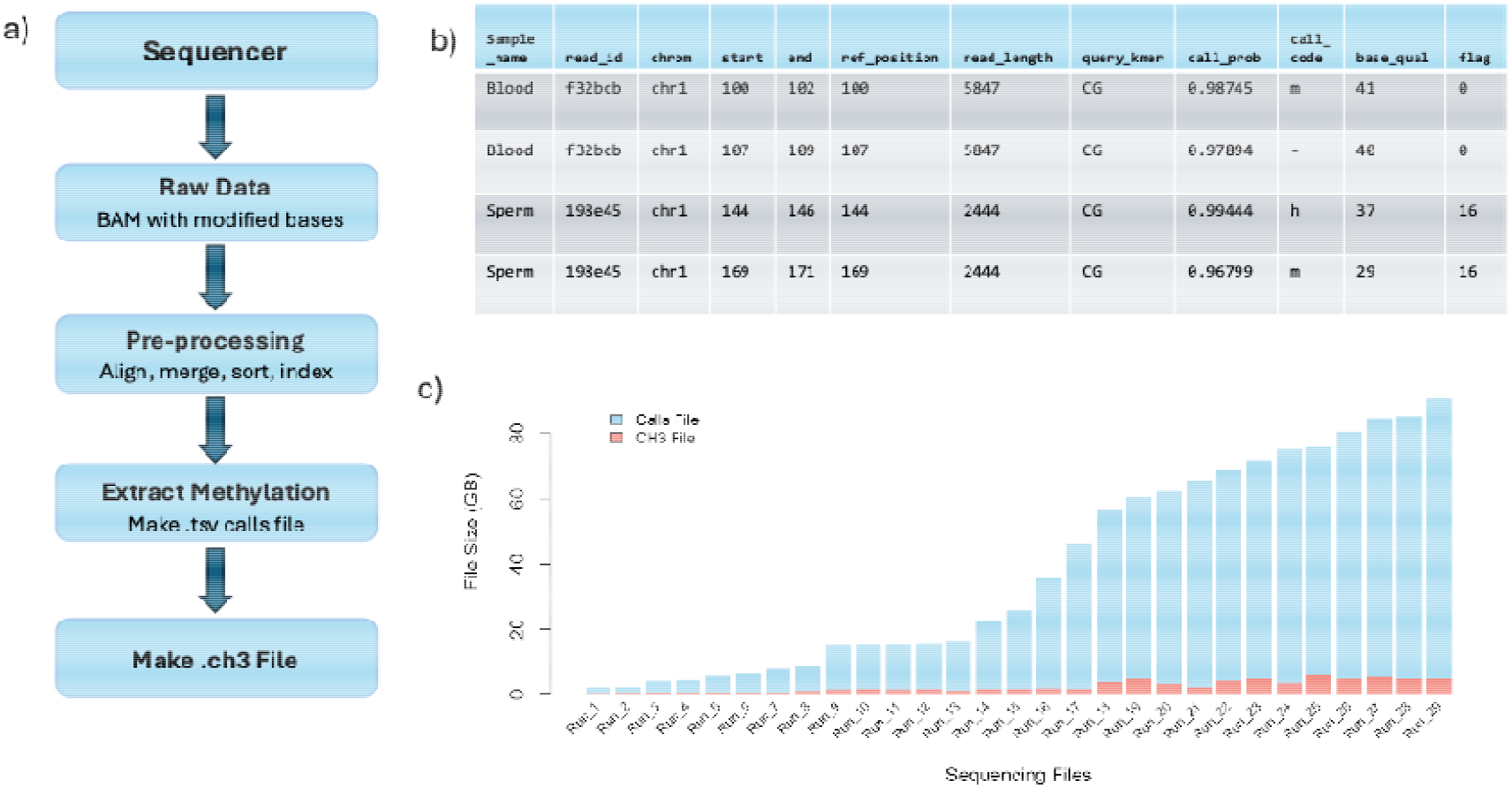
Specification for the CH3 file format. A) The general workflow used to generate a CH3 file. B) Sample CH3 file contents. C) Memory comparison between CH3 and modkit calls formats across 29 nanopore sequencing runs. Each bar represents the total file size in gigabytes (GB) for a single sequencing run, with modkit calls files shown in light blue and the corresponding CH3 files shown in red.

To address the limitations inherent in the standard methylation output format, we developed the custom CH3 file format (named after the chemical formula for a methyl group), a compact, efficient representation of base modification calls designed for efficient compression and retrieval. This format contains all essential nucleotide modification information, including the sample name, read ID, chromosome, 0-based reference position, read length, query k-mer, call probability, call code, base quality, and samtools flag value. The CH3 file specification is also extensible, allowing additional columns can be added to meet unique needs. Internally, the CH3 format leverages the Apache Parquet framework (https://parquet.apache.org/), enabling fast, column-based access and easy integration with many popular programming languages, including R, Python, C++, and Java. The Parquet format reduces file size by utilizing dictionary-based encoding for string columns and incorporates the zstd compression algorithm (https://github.com/facebook/zstd) to efficiently represent data within the file, allowing for fast access. In addition to the inherent efficiencies of the Parquet format, the CH3 file specification also relies on the samtools flag, which encodes alignment classification (unaligned, primary, secondary, or supplementary) and strand (+ or -) into a single integer instead of having multiple columns for this information (**Figure 1b**), further reducing its size. Together, these factors reduce the size of methylation data stored in TSV files to just 5–10% of their original size, ensuring that large-scale epigenomic datasets can be stored, transferred, and queried using minimal computational resources (**Figure 1c**).

### ModSeqR - An R Package for Analyzing Modified DNA

Current R packages for analyzing modified DNA have two major limitations. First, many are built around a bisulfite conversion paradigm. Thus, they do not support direct detection of modifications and are generally limited to methylation calls. However, third-generation sequencing platforms, such as the Nanopore platform, can directly detect a large number of modifications. Second, the amount of data generated from sequencing is extremely large. For example, there are approximately 30 million CpG dinucleotides in the human genome. At a sequencing depth of 10x, 300 million calls per sample, the data is orders of magnitude larger than the 900,000 data points generated by the Epic Array, and large enough to utilize the entire vector memory limit on machines with modest hardware specifications.

To address both of these issues, we have created the ModSeqR package (outlined in **Figure 2a**). This package implements a DuckDB database backend, enabling users to conduct large, complex analyses on a wide range of hardware^15^. The database is managed internally to maintain ease of use and familiarity for users accustomed to using R’s conventional data structures and paradigms (e.g. piping). The package also provides several built-in functions for working with the base call data to facilitate common workflows, including QC, data summarization by position, read, window, and region, and differential methylation analysis. An example pipeline with explanations for each function is shown in **Figure 2b-c**. Below, we outline the key steps of an analysis workflow and highlight the functionality and outputs of the package.

**Figure 2:**
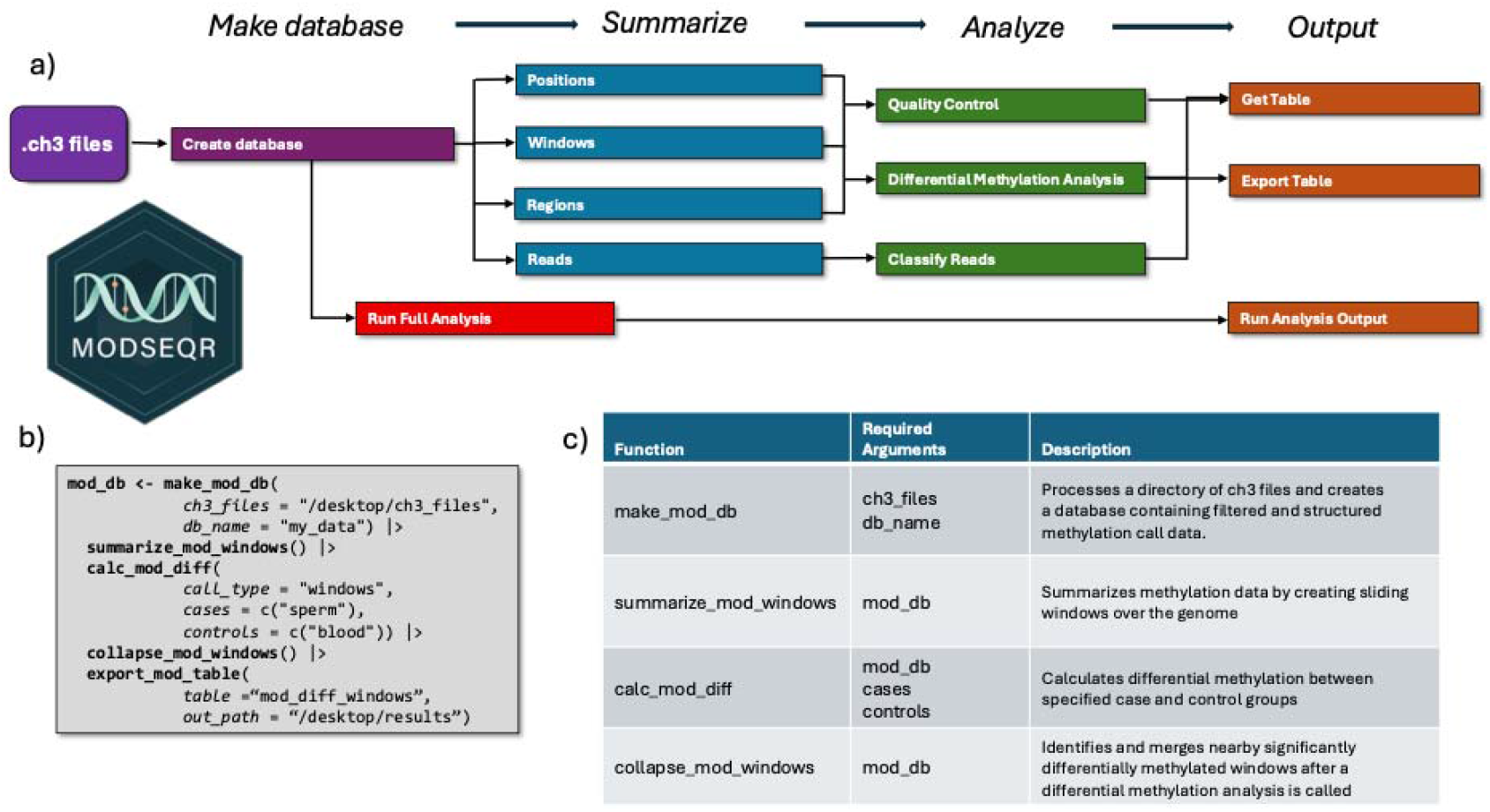
ModSeqR Core Design and Workflow. A) Schematic of possible ModSeqR workflows, starting from the input of CH3 files and progressing through database creation, summarization, analysis, and output. B) Example pipeline for a full differential methylation pipeline using R’s native pipe syntax. Pipeline begins with creating a database, then summarizes data using sliding windows, and performs a differential methylation analysis. Finally, significant windows are collapsed into merged regions, and the results are exported to a user-defined path. C) The core functions used in the example pipeline in Panel B, along with their required arguments and a brief description.

### Database Creation and Filtering

The first step in any ModSeqR pipeline is to create a database with all samples to be included in the analysis (**Figure 2**). The CH3 file format is designed to store the data from a single sample, while the database combines multiple samples into a single data structure, a calls table, for analysis (**Supplemental Figure 1**). The database is written to the local directory as “filename.ch3.db” and is updated as tables are created within it, allowing for the persistent storage of intermediate data structures and reducing the need to rerun calculations between sessions or to maintain large amounts of data in memory during multistep pipelines.

During this stage, the data can also be filtered for a variety of QC parameters, including minimum read length, minimum call probability, and minimum base quality. Additionally, users can choose to focus on specific types of modifications or combinations of several modifications. For example, users importing data from CH3 files containing methylation and hydroxymethylation can choose to use either modification, combine both into a single call, or retain all modifications. This level of customization empowers researchers to refine their workflows and explore epigenetic patterns in a way that aligns with their biological questions and data characteristics.

### Summarizing DNA methylation information over positions, pre-defined regions, or tiling windows

The modification calls stored in the database can be summarized in several ways (**Figure 2**). ModSeqR can summarize DNA methylation data across individual genomic positions, predefined regions (such as promoters, CpG islands, or introns), or sliding windows for further analysis. It also supports read-level summarization for mixture analysis (described below). These summaries can then be used for downstream analyses of these unique sites of interest. More detailed explanations of all functions within the package can be found in the ModSeqR vignette.

Users can export any resulting summary tables directly into their R environment for further exploration with the get_table() function. This includes data summarized by position, window, region, read, or a differential analysis. Example summarization table outputs using a synthetic dataset are shown in **Supplemental Figure 2**. Additionally, users can save the data table as a CSV file for external use by using export_table().

### Differential Methylation Analysis

A differential methylation analysis can be performed to identify statistically significant differences in DNA methylation levels at individual bases or genomic windows/regions between two different sample groups. This analysis is essential for detecting epigenetic changes that may play a role in gene regulation, development, or disease processes such as cancer and neurological disorders^16^. Differential methylation testing is performed using ModSeqR’s adaptive statistical framework (see Methods), which selects an appropriate hypothesis test based on replicate structure

#### Calibration of differential methylation testing

To evaluate the statistical calibration of ModSeqR’s differential methylation analysis, we generated synthetic nanopore modification datasets with known ground truth. Simulated datasets contained 1,000 CpG sites, of which 100 were assigned true methylation differences of ≥0.5 between case and control groups, while the remaining sites represented true nulls. Read coverage, per-read methylation probabilities, and replicate structure were simulated to reflect empirical nanopore sequencing data.

Differential methylation testing was performed using ModSeqR’s standard workflow, with p-values adjusted for multiple testing across all loci using the Benjamini– Hochberg procedure. At an adjusted p-value threshold of q < 0.05, ModSeqR identified 103 sites, of which 100 corresponded to true differentially methylated positions, yielding an empirical false discovery rate of 2.9% (**Supplemental Figure 3b**). Consistent with appropriate statistical calibration, p-values for true null sites were uniformly distributed prior to multiple testing correction and exhibited a conservative shift toward 1.0 following a Benjamini–Hochberg adjustment (**Supplemental Figure 3A**). Across a broad range of q-value thresholds, the empirical false discovery rate closely tracked or remained slightly below the nominal level (**Supplemental Figure 3c**), demonstrating effective control of false positives under realistic sequencing noise and replicate structure. Together, these results indicate that ModSeqR’s differential methylation analysis is well calibrated under realistic conditions, and that adjusted p-value thresholds provide reliable control of the false discovery rate.

**Figure 3:**
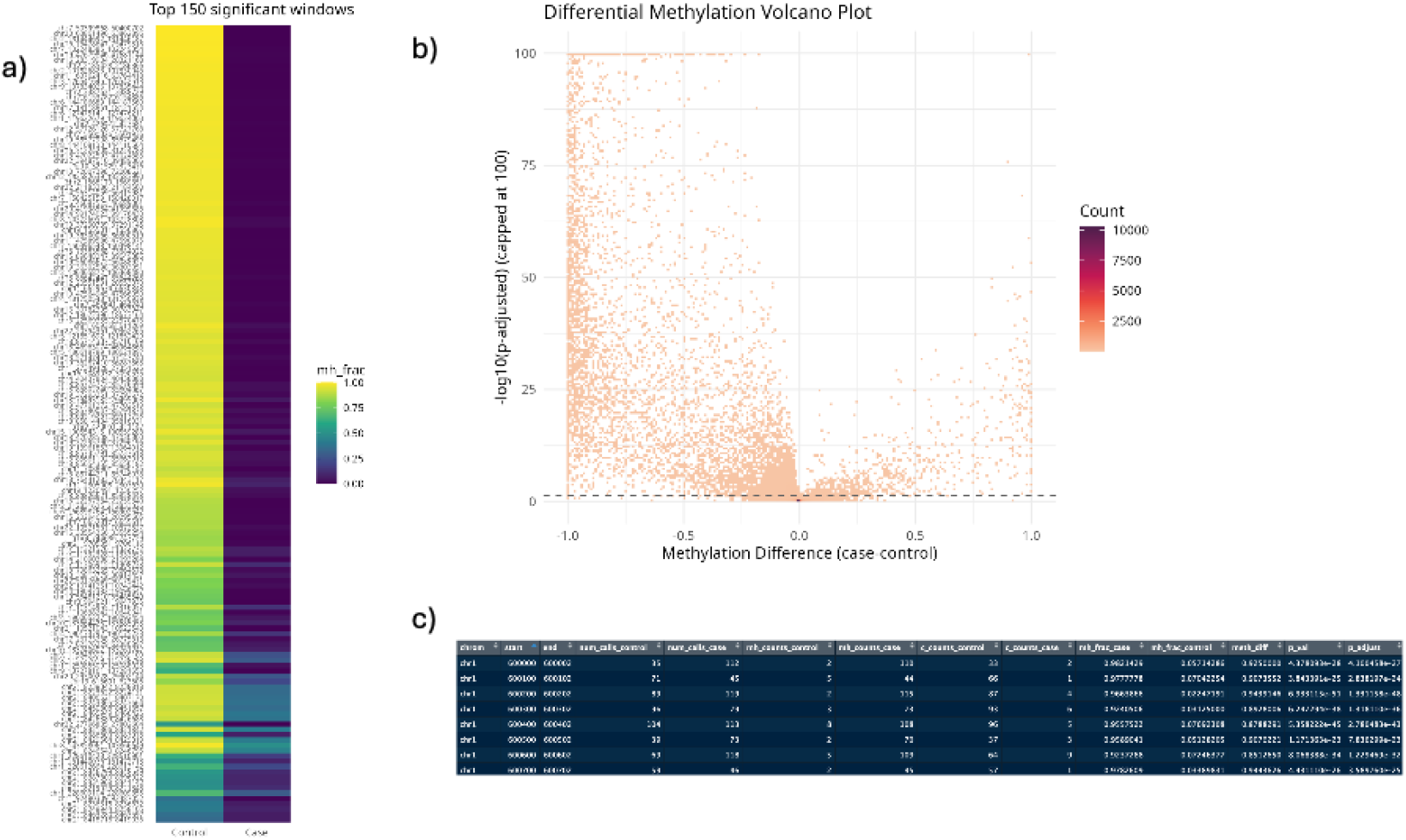
Example Output and Plots for Differential Expression Analysis. A) Heatmap showing the top 150 differentially methylated regions between sperm and blood. B) Example differential volcano plot generated with a custom ModSeqR function. Each square represents a genomic window/position. The x-axis represents the methylation-fraction difference between the sperm and blood samples, while the y-axis indicates the statistical significance (adjusted p-value). Density correlates with darker colors, and points near the center indicate regions with little or no difference C) An example R output table generated from the calc_mod_diff() function.

#### Outputs

The output of a differential methylation analysis can be directly plotted as a heatmap or a volcano plot, among others (**Figure 3a and b**). It can also be exported as a table to use with standard plotting functions in R **(Figure 3c)**.

### QC metrics: descriptive statistics, sample correlation, and PCA

In addition to its core analysis pipelines, ModSeqR provides a suite of tools for whole methylome characterization, including general quality control metrics, sample-level comparisons, and dimensionality reduction. Users can visually assess key characteristics of their methylation data using built-in plotting functions (**Figure 4**). Two primary visualization options are available: read coverage per base, which provides insight into sequencing depth and data reliability, and percent methylation per base, which reflects the underlying biological signal. These metrics are stored in the package’s core data structures and are easily accessible for customized visualizations.

**Figure 4:**
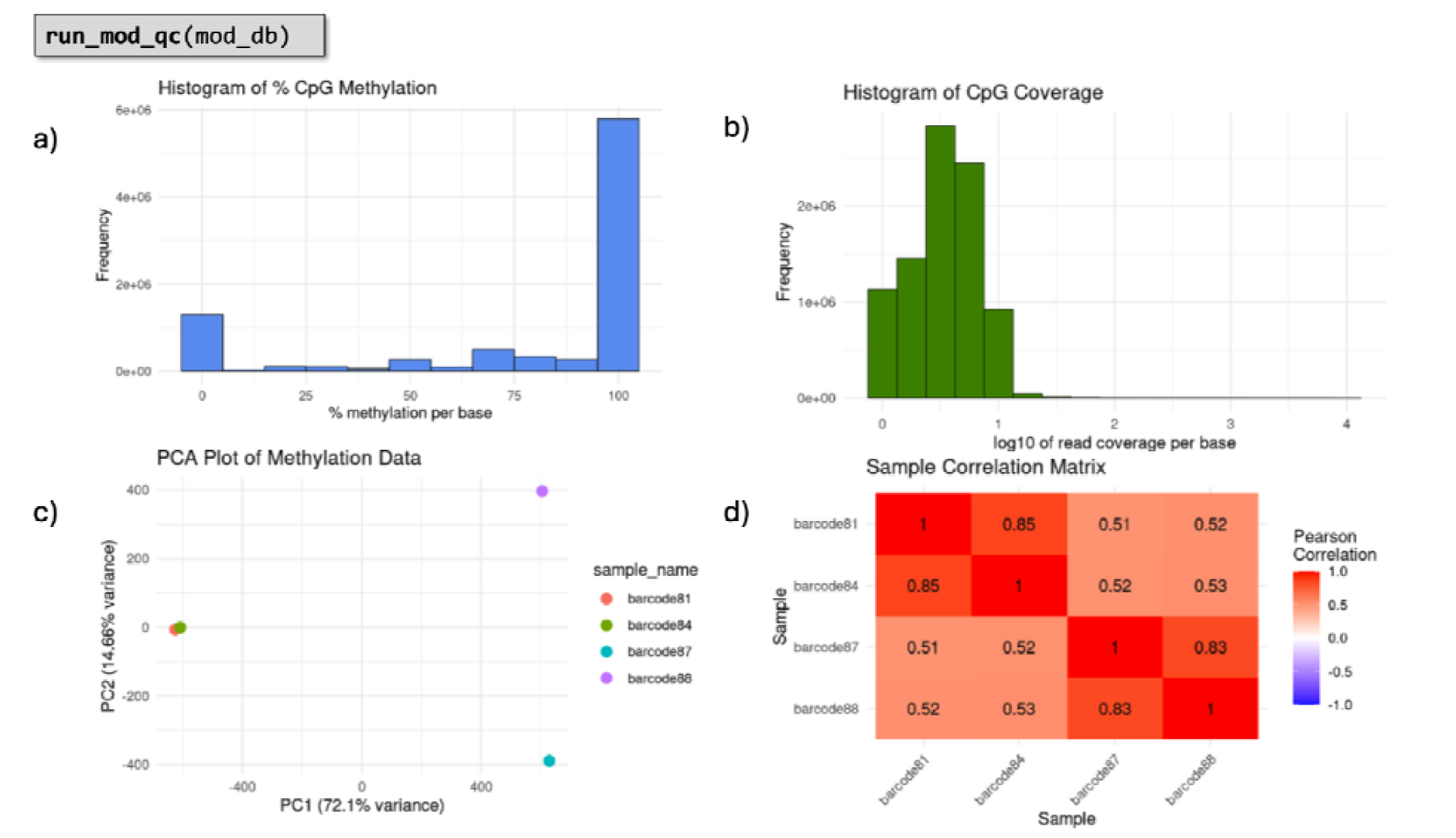
Descriptive Statistics and Quality Control Function Outputs. A) Percent methylation per base graph: This histogram shows the distribution of CpG methylation percentages across all bases for a sample. It provides an overview of the methylation landscape, allowing users to quickly assess whether methylation values are skewed, bimodal, or evenly distributed. B) Read coverage graph: This histogram displays the distribution of CpG read coverage (on a log10 scale) per base. It helps evaluate the depth and consistency of sequencing coverage, which is critical for assessing the reliability of methylation calls. C) PCA plot: The principal component analysis (PCA) plot reduces high-dimensional methylation data into two main components (PC1 and PC2) that capture the largest variance. This visualization helps identify clustering of samples and potential batch effects or outliers. D) Correlation plot: The sample correlation matrix helps visualize the pairwise Pearson correlation coefficients between samples based on methylation values. It quantifies the linear relationship between samples, helping assess similarity and reproducibility across experimental conditions.

To compare methylation profiles across samples, ModSeqR includes correlation plots based on the Pearson correlation coefficient, a measure of the linear relationship between two continuous variables. Additionally, a principal component analysis (PCA) function is provided to explore global patterns in methylation data and detect clustering among samples. PCA reduces the dimensionality of high-throughput data by projecting it into a smaller set of orthogonal components that capture the greatest variance, helping reveal biological trends or batch effects across the dataset.

### Performance

ModSeqR is designed to be computationally efficient and capable of analyzing large datasets in practical runtimes and on common hardware configurations. We tested the performance of several key analysis steps on various dataset sizes. The most computationally intense step within the analysis is summarization. The database is initially created with one row per methylation call. These calls are then summarized into positions, reads, regions, or sliding windows across all samples, which requires large amounts of database read and write calls. As shown in **Figure 5a**, the time required to conduct summarization depends on both the type of summarization and the dataset size. Importantly, however, the computation completes for all dataset sizes, even when they are larger than the available memory. Differential analysis also reveals a similar smooth increase over time without failure (**Figure 5b**). Finally, we compared run times between an analysis server with 250 GB of RAM and 96 CPU cores and a personal computer with 16 GB of RAM and 8 cores (**Figure 5c**). While run times were significantly longer on the personal computer, times were still reasonable, and all analyses were completed.

**Figure 5:**
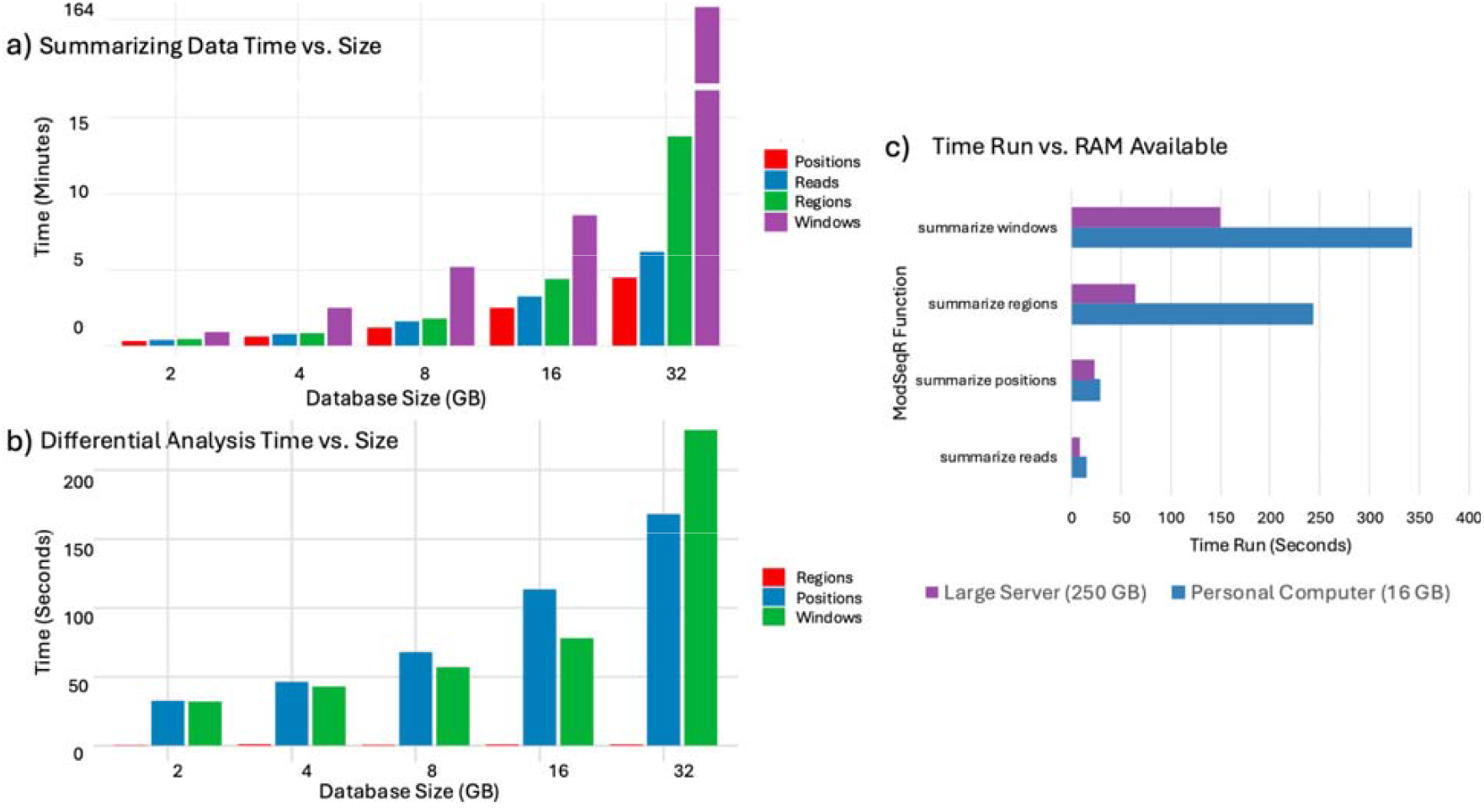
ModSeqR Performance on different hardware configurations. A) Summarizing Data Analysis Time vs. Size for different data collection functions. This includes summarizing methylation data by windows (100 bp step size), reads, genomic positions, or regions. B) Differential Analysis Time vs. Size. This figure depicts a differential analysis between three available options (regions, positions, and windows). These functions were run on a personal computer with 16GB of RAM. C) Time vs. RAM available for 4 functions on two different machines - one is a personal computer with 16GB of RAM available, while the other is a large server with 250GB of memory available. The dataset consisted of 4 samples of 5x coverage.

We also compared ModSeqR to four other software packages used to analyze differential methylation: diffONT, DSS, modkit, and PoreMeth2 (Table 1). We installed the latest version of each package, which we used to identify DMRs between one sperm sample of 5x coverage and one blood sample of 5x coverage. Note that both these samples were pulled from the dataset aforementioned. Each package was used to identify DMRs twice with the following settings: 16 GB RAM / 4 CPUs and 256 GB RAM / 16 CPUs (Table 2). With 16 GB RAM / 4 CPUs allotted, each package maxed out memory at 16 GB, and ModSeqR was by far the fastest analysis (9m 19s). With 256 GB RAM / 16 CPUs allotted, diffONT was the most efficient in memory usage (18.76 GB), but ModSeqR was again the fastest analysis (4m 8s).

**Table 1:** Feature Comparison of Five Differential Methylation Tools. A summary of each package’s key features, including sample QC methods, accepted modification types, and statistical analyses.

|  | Sequencing Type | Language | Sample QC | Modification Types | Multi-Modification Analysis | Differential Analyses | Read Classification | Data Plots |
| --- | --- | --- | --- | --- | --- | --- | --- | --- |
| ModSeqR | All | R | PCA on methylation fraction and coverage correlation | 5mC, 5hmC, 4mC, and 6mA (Any SAM/CHEBI code) | Yes | DMP, DMW, DMR | Yes | Coverage statistics, methylation histogram, methylation scatterplot, methylation volcano plot, PCA |
| diffONT | ONT | Python | None | 5mC or 5hmC | No | DMP, DMR | No | None |
| DSS | All | R | Minimum coverage parameters | 5mC, 5hmC, 4mC, and 6mA (Any SAM/CHEBI code) | No | DMP, DMR | No | Methylation line graph |
| modkit | ONT | Rust | Methylation fraction parameters | 5mC, 5hmC, 4mC, and 6mA (Any SAM/CHEBI code) | No | DMP, DMR | No | None |
| PoreMeth2 | ONT | Python | Shannon's Entropy parameters | 5mC or 5hmC | No | DMP, DMW, DMR | No | DMR $\Delta\beta$ / $\Delta S$ plot, DMR annotation plot |

**Table 2:**
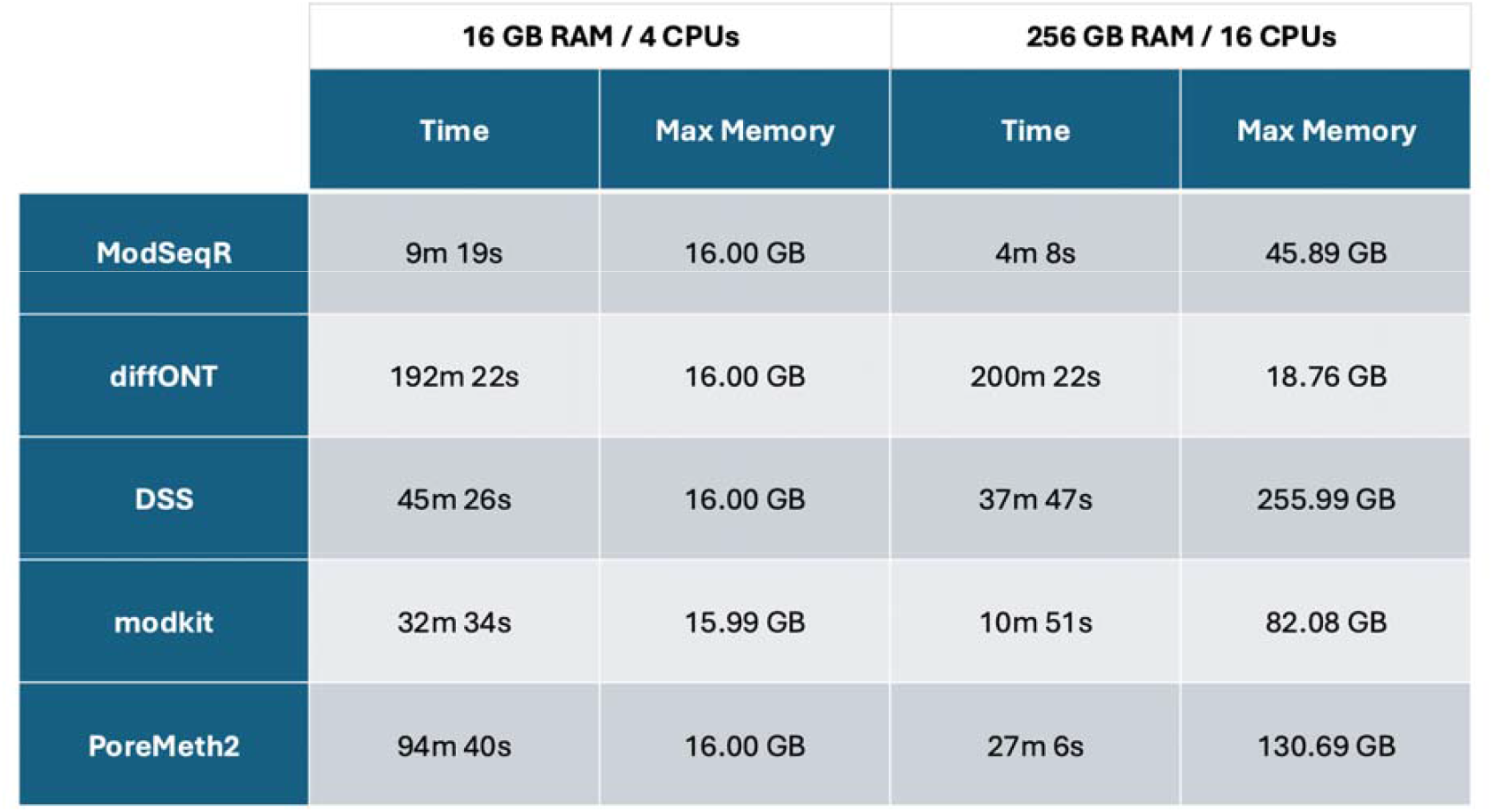
Performance Comparison of Five Differential Methylation Tools. Each package was used to identify DMRs between the same two samples using two different configurations: 16 GB RAM / 4 CPUs and 256 GB RAM / 16 CPUs. This table summarizes the wall-clock time and max memory usage for each analysis.

|  | 16 GB RAM / 4 CPUs |  | 256 GB RAM / 16 CPUs |  |
| --- | --- | --- | --- | --- |
|  | Time | Max Memory | Time | Max Memory |
| **ModSeqR** | 9m 19s | 16.00 GB | 4m 8s | 45.89 GB |
| **diffONT** | 192m 22s | 16.00 GB | 200m 22s | 18.76 GB |
| **DSS** | 45m 26s | 16.00 GB | 37m 47s | 255.99 GB |
| **modkit** | 32m 34s | 15.99 GB | 10m 51s | 82.08 GB |
| **PoreMeth2** | 94m 40s | 16.00 GB | 27m 6s | 130.69 GB |

## Discussion

Analyses of DNA methylation and other nucleotide modifications have enormous potential to provide novel biological insights into both natural processes and disease mechanisms. While such analyses have been conducted for over a decade, the recent development of direct, modified base calling on the NanoPore and PacBio platforms has significantly increased the genomic coverage of these assays. For example, the current Epic Array has approximately 900 thousand of the approximately 30 million CpG sites in the genome (approximately 3%). However, sequencing regularly provides information on over 28 million sites (93%). This represents a significant leap forward, providing valuable insights that were previously missed by other technologies.

Although the increased genomic coverage provided by third-generation sequencing is very welcome, it also results in a significant increase in the size of the data files produced, hindering analysis due to the requirement for large amounts of computational resources and specialized bioinformatics expertise to manage the data effectively. Here, we have presented two tools to address these issues: the CH3 file format and the ModSeqR R package. These tools, used together, will enable more users to perform modified nucleotide analysis and drive wider adoption across the field. Furthermore, our comparison of ModSeqR to four other differential methylation tools demonstrates that it significantly reduces the time and memory required, thereby enabling more users with limited computing resources to analyze large datasets more efficiently.

One of the primary challenges with large datasets lies in the cost and complexity of storing and transferring them—difficulties that will only intensify as data generation continues to grow. Thus, we created a file format specification, CH3, that addresses these problems by vastly decreasing the file size for modified base data. Analyses presented here show that CH3 typically reduces file sizes by 95%. These reductions come from a number of sources, including the removal of redundant and unnecessary data fields, efficient compression algorithms, and careful selection of data types. File access and read times are also improved by using column-based storage and internal indexing.

Importantly, the file specification is designed to be used with any modification code and allows users to add additional columns as needed to address specific use cases and future-proof the format. Thus, this file format will provide a strong foundation for future studies on nucleotide modifications.

We also present in this paper the first R package designed specifically for use with native, modified base calling. We have also incorporated a large number of functions to access and interact with the data, measure QC, and conduct differential methylation analyses on positions, sliding windows, annotated regions, or reads. As we look to the future, our team plans to further enhance ModSeqR’s analytical capabilities by integrating sophisticated machine learning algorithms, such as k-Nearest Neighbors (KNN) and regularized regression models, further advancing modified base analysis in various use cases^17^.

By dramatically reducing data storage requirements and offering an accessible, purpose-built R package, our work lowers key technical barriers that have limited the broader adoption of third-generation sequencing for epigenomic research. As the field continues to evolve, we anticipate that these tools will not only accelerate discovery but also inspire the development of new methodologies for understanding the biological roles of modified nucleotides in health and disease.

## Conclusions

The CH3 file format and the ModSeqR R package introduced in this study represent important advances for analyzing native DNA modifications using third-generation sequencing. By addressing key challenges in data size, accessibility, and analytical flexibility, these tools enable researchers to efficiently store, manage, and analyze high-throughput methylation data across a range of hardware configurations. The CH3 file format significantly reduces storage demands while preserving analytical utility, and ModSeqR provides a comprehensive and user-friendly R-based framework for differential methylation analysis, quality control, and visualization. Together, they remove critical technical barriers and empower broader use of native nucleotide modification data across biological research. As interest in epigenetic mechanisms continues to expand, these tools will serve as a foundational platform for scalable, accessible, and reproducible studies of DNA methylation and other base modifications in both health and disease contexts.

## Methods

### CH3 File Specification

To address the challenges of storing large-scale native base modification data, we developed the CH3 file format—a compressed, columnar format based on Apache Parquet. CH3 includes required fields such as read ID, chromosome, position, modification type, and probability, and supports additional optional metadata. File reduction is achieved via dictionary encoding and Zstandard compression, both natively implemented as part of the Parquet standard. The format also supports fast, column-based access and is compatible with ModSeqR and common data science tools (e.g., DuckDB, PyArrow). Full specification details, including field definitions and encoding rules, are available at the public GitHub repository https://github.com/Wasatch-Biolabs-Bfx/CH3. Additionally, a website has been developed for package explanation, a vignette, and function definitions at https://wasatch-biolabs-bfx.github.io/ModSeqR/.

### ModSeqR Implementation

ModSeqR is implemented in R and SQL and structured around modular functions that facilitate scalable, memory-efficient analysis of native base modification data. At its core, the package uses a DuckDB backend to support disk-based storage and efficient SQL-like querying of CH3-formatted data. Key functions are organized into workflow stages, including database creation (create_ch3_database()), read filtering (filter_ch3_reads()), summarization (summarize_ch3_positions(), summarize_ch3_windows(), summarize_ch3_regions()), and differential methylation analysis (calc_ch3_diff()). Additional tools provide sample correlation (calc_ch3_samplecor()), read classification (classify_ch3_reads()), quality control, and dimensionality reduction via PCA. Each function is optimized to perform reliably on systems with limited RAM and supports both single-sample and multi-sample analysis workflows. Extensive helper functions manage database connections, cleanup, and internal metadata. ModSeqR supports piping syntax and outputs tidy data frames, making it highly compatible with the broader tidyverse ecosystem. Full source code and documentation are available at: https://github.com/Wasatch-Biolabs-Bfx/ModSeqR.

#### Differential methylation analysis

ModSeqR provides multiple statistical frameworks for conducting differential methylation analysis, including logistic regression, Fisher’s exact test, a rapid implementation of Fisher’s exact test (r_fisher), and the Wilcoxon rank-sum test. Differential methylation can be assessed at individual genomic positions, sliding windows, or predefined genomic regions. For each locus, ModSeqR computes per-sample methylation fractions, from which group-level methylation summaries are derived. Effect sizes are reported as the difference in modification fraction between case and control groups, along with associated p-values.

For hypothesis testing, ModSeqR employs an adaptive test selection strategy based on the number of biological replicates in each group. When either the case or control group contains five or fewer samples, differential methylation significance is assessed using Fisher’s exact test applied to aggregated modified and unmodified read counts, an approach well suited for sparse data and low replicate numbers. When both case and control groups contain more than five samples, ModSeqR instead applies the Wilcoxon rank-sum test to compare distributions of per-sample methylation fractions between groups. This non-parametric test does not assume normality and leverages increased replicate numbers to provide improved robustness and sensitivity in analyses of larger cohorts. By dynamically selecting the appropriate statistical test based on sample size, ModSeqR ensures accurate and well-calibrated differential methylation inference across diverse experimental designs.

### Testing Data

We created a synthetic dataset to test the accuracy of our summarization functions and differential analysis. We hope this will also enable a fast workflow through all steps of the package. The positions are divided into deliberately chosen effect-size bands: eight groups with absolute methylation-fraction shifts of ±0.90, ±0.50, ±0.25 and ±0.10 (60 loci per band), and two mid-fraction bands (0.25–0.50 and 0.50–0.75; 120 loci each). We also included randomized noise for more realistic visuals. This dataset contains two CH3 files and is also publicly available for users to create a database and test the package under the inst/extdata/ directory in the GitHub repository.

#### Generation of Biological Data From Biological Data

DNA from sperm and blood samples were used to generate biologically relevant test data. 1 µg of fragmented isolated DNA was sent to Wasatch Biolabs (Heber City, Utah) for library preparation. Libraries were then sequenced at both Wasatch Biolabs and Brigham Young University.

All sequencing was conducted using ONT PromethION flow cells (FLO-PRO114) according to the manufacturer’s instructions. Sequencing, basecalling, and demultiplexing were performed using MinKNOW software with live high-accuracy basecalling selected and with 5mC and 5hmC modifications in the CpG context turned on. Reads with a Q score less than 10 were marked as “failed” and were removed from subsequent analysis.

### Performance Benchmarking

Performance testing was conducted on two markedly different hardware configurations to assess scalability and resource utilization. The first environment was a MacBook Pro equipped with an M3 processor and 16 GB of RAM, representing a typical end-user system. The second environment was a high-performance server with 250 GB of available memory and an AMD Threadripper processor with 96 cores, providing substantially greater computational capacity.

To compare the performance of ModSeqR to other differential methylation tools, we utilized an HPC cluster with Slurm job scheduling. Using Apptainer, we deployed the latest version of each package in separate containers, along with their respectively required dependencies. Each analysis was submitted with either 16 GB RAM / 4 CPUs or 256 GB RAM / 16 CPUs via Slurm resource allocations. Wall-clock time and max memory usage were automatically recorded for each submission.

## Declarations

## Software Availability and Requirements

**Project name:** ModSeqR

**Project home page:** https://github.com/Wasatch-Biolabs-Bfx/ModSeqR

**Operating system(s):** Platform independent (works on Windows, macOS, and Linux wherever R is available).

**Programming language:** R

**Other requirements:**R package dependencies include: arrow, DBI, dplyr, dbplyr, duckdb, duckplyr, ggplot2, progress, tidyr, utils, readr (Imports). Optional for vignettes/docs: knitr, rmarkdown (Suggests).

**License:**Personal and Internal Research License (v1.1) Based on PolyForm Noncommercial License 1.0.0, with an Additional Grant of Rights (Internal Research Use Exception) provided for internal research use by commercial entities.

Any restrictions to use by non-academics: Internal evaluation and research only.

## Ethics approval and consent to participate

All human-derived materials were collected in accordance with relevant institutional guidelines and regulations and in compliance with the Declaration of Helsinki. Sperm samples were obtained under a protocol reviewed by the Brigham Young University Institutional Review Board (IRB) and determined to be exempt from IRB oversight (Protocol #A2019-349). Menstrual blood samples were collected under a protocol approved by the Brigham Young University IRB (Protocol #IRB2022-245). Informed consent was obtained from all participants where applicable.

## Consent for publication

Not applicable

## Availability of data and materials

The dataset analyzed during the current study is available in the ModSeqR repository through the Wasatch-Biolabs-Bfx organization, at https://github.com/Wasatch-Biolabs-Bfx/ModSeqR. Documentation and a package vignette is publicly available on our package website at https://wasatch-biolabs-bfx.github.io/ModSeqR/. Sequencing data is available in the NCBI SRA database under project number PRJNA1311512.

## Authors’ contributions

HZ and JH wrote the code for the ModSeqR package, as well as writing this manuscript. RM, JM, and AJ assisted in developing the package and writing this paper. IS assisted in writing and figure generation. ES assisted in writing and editing the manuscript. TJ helped oversee the project and provided guidance for methylation analysis standards.

