## Supplemental Figure 1 for "ModSeqR: An R package for efficient analysis of modified nucleotide data" (from another version of this preprint: version 1, 2025-11-12)

a) Calls table example

| sample_name | read_id | chrom | start | end | read_position | ref_position | read_length | query_kmer |
|---|---|---|---|---|---|---|---|---|
| case | case_read_1_0 | chr1 | 600000 | 600002 | 1 | 600000 | 1000 | CG |
| case | case_read_1_1 | chr1 | 600000 | 600002 | 6 | 600000 | 1000 | CG |
| case | case_read_1_2 | chr1 | 600000 | 600002 | 11 | 600000 | 1000 | CG |
| case | case_read_1_3 | chr1 | 600000 | 600002 | 16 | 600000 | 1000 | CG |
| case | case_read_1_4 | chr1 | 600000 | 600002 | 21 | 600000 | 1000 | CG |
| case | case_read_1_5 | chr1 | 600000 | 600002 | 26 | 600000 | 1000 | CG |
| case | case_read_1_6 | chr1 | 600000 | 600002 | 31 | 600000 | 1000 | CG |
| case | case_read_1_7 | chr1 | 600000 | 600002 | 36 | 600000 | 1000 | CG |
| case | case_read_1_8 | chr1 | 600000 | 600002 | 41 | 600000 | 1000 | CG |
| case | case_read_1_9 | chr1 | 600000 | 600002 | 46 | 600000 | 1000 | CG |
| case | case_read_1_10 | chr1 | 600000 | 600002 | 51 | 600000 | 1000 | CG |
| case | case_read_1_11 | chr1 | 600000 | 600002 | 56 | 600000 | 1000 | CG |
| case | case_read_1_12 | chr1 | 600000 | 600002 | 61 | 600000 | 1000 | CG |
| case | case_read_1_13 | chr1 | 600000 | 600002 | 66 | 600000 | 1000 | CG |
| case | case_read_1_14 | chr1 | 600000 | 600002 | 71 | 600000 | 1000 | CG |
| case | case_read_1_15 | chr1 | 600000 | 600002 | 76 | 600000 | 1000 | CG |
| case | case_read_1_16 | chr1 | 600000 | 600002 | 81 | 600000 | 1000 | CG |
| case | case_read_1_17 | chr1 | 600000 | 600002 | 86 | 600000 | 1000 | CG |
