## Supplemental Figure 2 for "ModSeqR: An R package for efficient analysis of modified nucleotide data" (from another version of this preprint: version 1, 2025-11-12)

### a) Positions table

| sample_name | chrom | start | end | num_calls | c_counts | m_counts | h_counts | mh_counts | m_frac | h_frac | mh_frac |
|---|---|---|---|---|---|---|---|---|---|---|---|
| control | chr1 | 600000 | 600002 | 35 | 33 | 1 | 1 | 2 | 0.028571429 | 0.028571429 | 0.05714286 |
| case | chr1 | 600000 | 600002 | 112 | 2 | 55 | 55 | 110 | 0.491071429 | 0.491071429 | 0.98214286 |
| case | chr1 | 600100 | 600102 | 45 | 1 | 22 | 22 | 44 | 0.488888889 | 0.488888889 | 0.97777778 |
| control | chr1 | 600100 | 600102 | 71 | 66 | 3 | 2 | 5 | 0.042253521 | 0.028169014 | 0.07042254 |
| case | chr1 | 600200 | 600202 | 119 | 4 | 58 | 57 | 115 | 0.487394958 | 0.478991597 | 0.96638655 |

### b) Windows table

| sample_name | chrom | start | end | num_CpGs | num_calls | c_counts | m_counts | h_counts | mh_counts | m_frac | h_frac | mh_frac |
|---|---|---|---|---|---|---|---|---|---|---|---|---|
| case | chr1 | 599001 | 600000 | 1 | 112 | 2 | 55 | 55 | 110 | 55.000000 | 55.000000 | 110.000000 |
| control | chr1 | 599001 | 600000 | 1 | 35 | 33 | 1 | 1 | 2 | 1.000000 | 1.000000 | 2.000000 |
| control | chr1 | 599011 | 600010 | 1 | 35 | 33 | 1 | 1 | 2 | 1.000000 | 1.000000 | 2.000000 |
| case | chr1 | 599011 | 600010 | 1 | 112 | 2 | 55 | 55 | 110 | 55.000000 | 55.000000 | 110.000000 |
| case | chr1 | 599021 | 600020 | 1 | 112 | 2 | 55 | 55 | 110 | 55.000000 | 55.000000 | 110.000000 |

### c) Regions table

| sample_name | region_name | chrom | start | end | num_CpGs | num_calls | c_counts | m_counts | h_counts | mh_counts | m_frac | h_frac | mh_frac |
|---|---|---|---|---|---|---|---|---|---|---|---|---|---|
| case | chr1_1231621_1233605 | chr1 | 1231621 | 1233605 | 20 | 1574 | 185 | 699 | 690 | 1389 | 0.44409149 | 0.43837357 | 0.88246506 |
| control | chr1_1231621_1233605 | chr1 | 1231621 | 1233605 | 20 | 1238 | 1072 | 89 | 77 | 166 | 0.07189015 | 0.06219709 | 0.13408724 |
| case | chr1_1242505_1242827 | chr1 | 1242505 | 1242827 | 3 | 221 | 196 | 13 | 12 | 25 | 0.05882353 | 0.05429864 | 0.11312217 |
| control | chr1_1242505_1242827 | chr1 | 1242505 | 1242827 | 3 | 243 | 22 | 111 | 110 | 221 | 0.45679012 | 0.45267490 | 0.90946502 |
| control | chr1_1246376_1247090 | chr1 | 1246376 | 1247090 | 7 | 474 | 71 | 203 | 200 | 403 | 0.42827004 | 0.42194093 | 0.85021097 |
