## Supplemental Figure 3 for "ModSeqR: An R package for efficient analysis of modified nucleotide data" (from another version of this preprint: version 1, 2025-11-12)

### a) Differential methylation analysis output table

| chrom | start | end | num_calls_control | num_calls_case | mh_counts_control | mh_counts_case | c_counts_control | c_counts_case | mh_frac_case | mh_frac_control | meth_diff | p_val | p_adjust |
|---|---|---|---|---|---|---|---|---|---|---|---|---|---|
| chr1 | 600000 | 600002 | 35 | 112 | 2 | 110 | 33 | 2 | 0.9821429 | 0.05714286 | 0.9250000 | 4.378093e-28 | 4.100458e-27 |
| chr1 | 600100 | 600102 | 71 | 45 | 5 | 44 | 66 | 1 | 0.9777778 | 0.07042254 | 0.9073552 | 3.843391e-25 | 2.838197e-24 |
| chr1 | 600200 | 600202 | 89 | 119 | 2 | 115 | 87 | 4 | 0.9663866 | 0.02247191 | 0.9439146 | 6.933115e-51 | 1.331158e-48 |
| chr1 | 600300 | 600302 | 96 | 79 | 3 | 73 | 93 | 6 | 0.9240506 | 0.03125000 | 0.8928006 | 6.247294e-38 | 1.318110e-36 |
| chr1 | 600400 | 600402 | 104 | 113 | 8 | 108 | 96 | 5 | 0.9557522 | 0.07692308 | 0.8788291 | 5.358222e-45 | 2.780483e-43 |
| chr1 | 600500 | 600502 | 39 | 73 | 2 | 70 | 37 | 3 | 0.9589041 | 0.05128205 | 0.9076221 | 1.171363e-23 | 7.836299e-23 |
| chr1 | 600600 | 600602 | 69 | 118 | 5 | 109 | 64 | 9 | 0.9237288 | 0.07246377 | 0.8512650 | 8.068388e-34 | 1.229469e-32 |
| chr1 | 600700 | 600702 | 59 | 46 | 2 | 45 | 57 | 1 | 0.9782609 | 0.03389831 | 0.9443626 | 4.431110e-26 | 3.589760e-25 |
| chr1 | 600800 | 600802 | 70 | 32 | 3 | 31 | 67 | 1 | 0.9687500 | 0.04285714 | 0.9258929 | 1.337667e-21 | 7.951456e-21 |
| chr1 | 600900 | 600902 | 118 | 105 | 4 | 98 | 114 | 7 | 0.9333333 | 0.03389831 | 0.8994350 | 5.466603e-49 | 4.563425e-47 |
| chr1 | 601000 | 601002 | 105 | 101 | 5 | 93 | 100 | 8 | 0.9207921 | 0.04761905 | 0.8731730 | 4.388287e-42 | 1.504555e-40 |
| chr1 | 601100 | 601102 | 60 | 88 | 3 | 85 | 57 | 3 | 0.9659091 | 0.05000000 | 0.9159091 | 2.275396e-33 | 3.334932e-32 |
| chr1 | 601200 | 601202 | 57 | 91 | 2 | 84 | 55 | 7 | 0.9230769 | 0.03508772 | 0.8879892 | 3.875500e-30 | 4.351438e-29 |
| chr1 | 601300 | 601302 | 46 | 96 | 3 | 92 | 43 | 4 | 0.9583333 | 0.06521739 | 0.8931159 | 5.027580e-28 | 4.640843e-27 |
| chr1 | 601400 | 601402 | 101 | 48 | 7 | 45 | 94 | 3 | 0.9375000 | 0.06930693 | 0.8681931 | 6.118121e-27 | 5.388437e-26 |
| chr1 | 601500 | 601502 | 89 | 68 | 6 | 63 | 83 | 5 | 0.9264706 | 0.06741573 | 0.8590549 | 1.657531e-30 | 1.928763e-29 |
| chr1 | 601600 | 601602 | 30 | 33 | 1 | 32 | 29 | 1 | 0.9696970 | 0.03333333 | 0.9363636 | 1.152446e-15 | 5.449990e-15 |
| chr1 | 601700 | 601702 | 73 | 61 | 4 | 60 | 69 | 1 | 0.9836066 | 0.05479452 | 0.9288120 | 5.073064e-32 | 6.581272e-31 |
| chr1 | 601800 | 601802 | 59 | 79 | 4 | 76 | 55 | 3 | 0.9620253 | 0.06779661 | 0.8942287 | 8.801136e-30 | 9.493360e-29 |
| chr1 | 601900 | 601902 | 83 | 39 | 2 | 37 | 81 | 2 | 0.9487179 | 0.02409639 | 0.9246216 | 2.060368e-26 | 1.705132e-25 |
| chr1 | 602000 | 602002 | 36 | 31 | 2 | 29 | 34 | 2 | 0.9354839 | 0.05555556 | 0.8799283 | 2.475716e-14 | 1.085245e-13 |
| chr1 | 602100 | 602102 | 82 | 51 | 3 | 47 | 79 | 4 | 0.9215686 | 0.03658537 | 0.8849833 | 1.796815e-27 | 1.635017e-26 |
